# Latent Representations of Early Brain Development: A Multivariate Normative Model of Brain Structure and Behaviour

**DOI:** 10.1101/2025.08.14.668769

**Authors:** Mariam Zabihi, Francesca Biondo, Jonathan O’Muircheartaigh, Thomas Wolfers, Sean Deoni, Andre Marquand, Muriel M.K. Bruchhage, James H. Cole

## Abstract

Individual variation in neurodevelopment plays a central role in shaping cognitive abilities and behavioural profiles, influencing both typical functioning and risk for neurodevelopmental conditions. While much research has focused on characterising trajectories of brain structure changes during development, this typically entails assessing brain regions individually, overlooking the multivariate nature of neuroimaging data. In this study, we trained an autoencoder to map latent representations of children’s brain development using T1-weighted MRI scans from a paediatric cohort (n = 564, 55% male, mean age 4.90 years, age range [0.11,15.38]). This approach enabled us to establish multivariate normative models of brain structure, providing a more comprehensive framework for understanding neurodevelopmental variation.

The latent space representation from this model effectively captured demographic variables (age and sex), while preserving both global and local structural features. The model accurately reconstructed the data, having mean reconstruction of *0.04 ± 0.01, whilst also capturing demographic features* with classification accuracy for sex of *84% ± 4%*, and a mean absolute error of *0.79 ± 0.06 years* for age prediction, highlighting its sensitivity to developmental changes. Further, we validated the approach using correlation analysis to show that deviations from the latent norms were significantly associated with multiple cognitive and behavioural measures, suggesting that variations in brain structure may reflect individual differences in neurodevelopment. Finally, we generated reference brain images that represent typical development and used them to visualise structural differences in individuals who deviate from this normative pattern. Our findings demonstrate that semi-supervised autoencoders, combined with multivariate normative modelling, offer a framework for characterizing neurodevelopmental trajectories. This approach can identify meaningful deviations associated with cognition and behaviour and has potential future applications across the lifespan.

## Introduction

The human brain undergoes a remarkable metamorphosis throughout life, with the most dramatic changes occurring during early childhood. This period of rapid development is characterized by intricate structural and functional alterations, which are heavily influenced by critical periods of neuroplasticity[1,2]. These windows of heightened brain malleability enable the formation of key neural circuits, laying the foundation for cognitive abilities, language, motor functions, as well as behavioural patterns. The interplay between genetic factors and environmental inputs, such as early experiences and social interactions, further shapes these developmental trajectories, contributing to individual differences in cognition and behaviour. Mapping this complex developmental process and capturing its inherent variability presents a formidable challenge for neuroscientists, with far-reaching implications for our understanding of cognitive development and associated disorders [3–5].

A growing body of research has employed MRI to study neurodevelopmental trajectories in children and infants, revealing significant insights into both typical and atypical brain development [6–9]. MRI-based studies have demonstrated that the brain’s volume, cortical thickness, and white matter organization undergo rapid transformations during the first years of life, with distinct patterns of maturation observed across different regions [10]. These findings not only advance our understanding of the brain’s structural evolution but also underline the critical importance of sensitive periods in shaping language, executive function, and social cognition. Identifying deviations during these key periods can offer early markers for neurodevelopmental disorders such as autism spectrum disorder (ASD) and attention-deficit/hyperactivity disorder (ADHD), facilitating early interventions[11,12].

In recent years, normative modelling has emerged as a powerful tool for investigating neurodevelopmental trajectories [13–19]. Drawing parallels with paediatric growth charts, this approach maps individual brain data against a reference population, effectively capturing variability and delineating what may be considered a ‘common’ range of development. By enabling the identification and quantification of deviations from typical development, normative modelling offers valuable insights into brain-behaviour relationships and potential indicators of cognitive or behavioural challenges.

Despite their applications, traditional normative models have limitations. These approaches often rely on predetermined, manually selected features, such as regions of interest (ROIs) or image-derived phenotypes (IDPs) [16,19–22]. While these features can yield valuable insights, they may not fully capture the intricate complexity of the brain, particularly in the context of the rapid and multifaceted changes occurring throughout development. On the other hand, applying normative models directly to raw inputs, such as voxel-wise or vertex-wise data[17,23], provide limited insight into coordinated patterns of brain-wide maturation.

To address these limitations, we propose an autoencoder framework that learns low-dimensional, multivariate representations of paediatric brain structure from high-dimensional neuroimaging data [24]. By reducing the data’s dimensionality into a small number of latent variables, the model captures coordinated axes of variation that reflect biologically meaningful developmental processes. This compact latent space supports normative modelling in a way that is both scalable and interpretable, enabling the detection of individual deviations while preserving global and local structural information. In contrast to voxel-wise methods, which produce spatially dense but unstructured deviation maps, our latent representation provides a parsimonious summary of brain development that can be related directly to cognitive and behavioural measures. Our methodology focuses on balancing the trade-off between model complexity and interpretability, ensuring that the latent features extracted are not only predictive but also biologically or behaviourally relevant. By providing a proof-of-principle, we aim to contribute to a more complete understanding of neurodevelopmental trajectories, paving the way for better identification of early markers of educational outcomes and cognitive or behavioural disorders.

## Methods and Materials

### Imaging Data

Brain imaging data were from the **BAMBAM** (Brown University Assessment of Myelination and Behavioural Development Across Maturation) study, which focuses on neurotypical brain and cognitive development [25]. This study, conducted at Brown University in Providence, RI, USA, was designed as an accelerated-longitudinal investigation of a large community cohort of healthy children. About half of the participants were enrolled between 2-8 months of age, while the remainder joined between 2-4 years of age [25]. Here, we used T1-weighted MRI scans from 564 (251 females, 313 males) participants, which included a total of N=1027 scans (584 male and 443 female scans, age = 4.90 ± 3.67 [0.11, 15.38]). T1-weighted anatomical images were acquired using a 3T Siemens Trio scanner equipped with a 12-channel head RF array. The imaging sequence employed a magnetization-prepared rapid acquisition gradient echo (MPRAGE) protocol, producing isotropic voxel data with an initial resolution of 1.2 × 1.2 × 1.2 mm^3^, later resampled to 0.9 × 0.9 × 0.9 mm^3^. Sequence parameters included: echo time (TE) = 6.9 ms, repetition time (TR) = 16 ms, inversion time = 950 ms, flip angle = 15°, and bandwidth = 450 Hz/pixel. To ensure consistent voxel volume and spatial resolution across all ages, the acquisition matrix and field of view were adjusted based on each child’s head size [9]. Cross-sectional processing allowed us to capture a broad representation of developmental stages while ensuring sufficient statistical power for our machine learning models.

To mitigate computational load while preserving structural details, we downsampled the scans from an initial resolution of 1 mm^3^ to 3 mm^3^ voxel size. This downsampling step, coupled with a tight cropping around the whole brain, resulted in image dimensions of 64×72×72. Furthermore, all images were normalized to have zero mean and unit variance across voxels, a preprocessing step designed to standardize the input data and facilitate more effective learning by the autoencoder.

For model validation, we employed five-fold cross-validation. The dataset was randomly partitioned into five mutually exclusive subsets, ensuring that each fold contained a distinct set of participants with no overlap across folds. This cross-validation procedure was designed to improve generalizability of the results and minimize overfitting, a common challenge in machine learning models trained on high-dimensional neuroimaging data.

### Brain features

To explore how conventional brain features relate to the learned latent representations, we included total brain volume, white matter volume(WMV), cortical grey matter volume(GMV), and subcortical grey matter volume(sGMV) as brain features, based on estimates previously derived from the BAMBAM dataset [26]. T1-weighted MRI scans were first registered to age-specific templates and then aligned to a common reference space. Brain regions were labelled using the Mindboggle OASIS atlas [27], allowing volumes of cortical and subcortical grey matter to be calculated by summing the relevant regions. White matter volume was estimated from tissue segmentations and adjusted by subtracting grey matter volumes from the overall brain tissue estimate. Total volume was defined as the sum of white matter, cortical grey matter, and subcortical grey matter volumes.

### Cognitive and Behavioural Data

In addition to imaging data, we integrated 139 cognitive and behavioural measures derived from widely used assessment scales. These were the younger and older children Child Behaviour Checklist (CBCL)[28], the Mullen Scales of Early Learning [29], the Wechsler Preschool and Primary Scale of Intelligence (WPPSI-IV) [30], the NEPSY-II (A Developmental Neuropsychological Assessment) [31], the Behaviour Rating Inventory of Executive Function (BRIEF)[32], the Comprehensive Test of Phonological Processing (CTOPP-2) [33], and the Beery-Buktenica Developmental Test of Visual-Motor Integration (Beery-VMI) [34]. These instruments assess a wide range of clinical, cognitive, and behavioural domains, including mental health, executive function, phonological processing, and visual-motor integration.

Given the strong developmental dependency of many of these measures, we used age-corrected scores where available, to ensure that observed associations reflect individual variability beyond expected age effects.

By leveraging this diverse array of assessments, we aimed to establish a comprehensive relationship between latent brain representations and behavioural measures. This multimodal integration allows us to explore not just structural correlates of neurodevelopment, but also how these structural variations might predict cognitive and behavioural performance. Importantly, the cognitive and behavioural data provide external validity to the model by anchoring its latent representations to meaningful real-world measures, thus enhancing the interpretability and translational potential of the findings.

### Normative Latent space

We employed a 3-dimensional conditional autoencoder to learn latent representations of structural MRI data, based on our previous work (Zabihi et al., 2024). The model was designed to jointly minimise reconstruction error and prediction loss for demographic variables (age and sex), encouraging the latent space to encode developmentally meaningful features.

The network architecture consists of three convolutional layers in both the encoder and decoder, connected via a fully connected latent layer of size 10. All convolutional layers used 3×3×3 kernels to capture both local and global image features. To promote sparsity and interpretability, we applied L1 regularization to the latent layer and removed bias terms from all layers. Weights were initialized using Xavier initialization [35]. The model was trained for up to 1,000 epochs with early stopping (patience = 10), using the Adam optimizer[36] [28] with an exponentially decaying learning rate (initial rate = 0.01, final rate = 0.0001), and a mini-batch size of 32.

Next, we applied normative modelling, using hierarchical Bayesian regression (HBR) [37] to the 10-dimensional latent representations extracted from autoencoder. We used a Gaussian likelihood in the HBR model for consistency across latents. While this assumption may not capture all non-Gaussian properties of the latent distributions, it provides a well-established and interpretable baseline. Here, we modelled age as a continuous covariate and sex as a batch effect to predict the “normative latent space”. By modelling brain data in a normative framework, we removed both linear and non-linear associations of age and sex, effectively defining a ‘reference’ brain state for each demographic group. This approach is particularly suited for neurodevelopmental studies, where it is critical to identify deviations from typical developmental trajectories. We then calculated a ‘latent index’ for each individual, representing their deviation from the normative latent space, using a subject-specific Z-score (as described in [17]). To connect these latent representations back to cognitive and behavioural measures, we applied multiple regression to show the linear associations between the latent index and 139 assessments from the BAMBAM dataset.

Finally, to generate normative brain images, we passed the predicted latent vectors from the HBR model into the decoder of the trained autoencoder. This enabled us to reconstruct synthetic brain scans representing the typical structural pattern expected at a given age and sex. Similarly, to visualise inter-individual variability, we sampled latent vectors at different centiles (e.g., 0.05, 0.95) and decoded them into corresponding MRI images. Here, we examined developmental changes by subtracting reconstructed images across age pairs (e.g., 54 months minus 4 months), yielding voxel-wise intensity difference maps that reflect typical maturational patterns over time.

## Results

We evaluated the performance of our semi-supervised autoencoder using three primary metrics, with results averaged across five cross-validation folds. First, we assessed the model’s ability to reconstruct the original brain imaging data from its latent representation. The image reconstruction error was **0.04±0.01**, demonstrating that the autoencoder effectively compresses and reconstructs the brain MRI data with minimal information loss. The corresponding **R**^**2**^ was **0.76±0.13**, indicating that the model explains a substantial proportion of the variance in the input data.

Next, we evaluated the model’s ability to predict demographic variables. Our model achieved an accuracy of **84%±4%** in sex prediction, indicating that it could effectively differentiate between male and female brain structures. For age prediction, the model had a mean absolute error (MAE) of **0.79±0.06** years and out of sample **R**^**2**^**= 0.91±0.1** underscoring its sensitivity to age-related variations in structural MRI data.

The correlations of the 10 latent variables with brain features including total white matter volume, total grey matter volume, subcortical grey matter volume, and total brain volume are shown in Figure 1.A. Figure 1.B shows the normative model of each latent variable by age. This normative model acts as a reference framework, helping to identify deviations in brain structure across developmental stages in the reference model. These curves define the centiles (e.g., 0.05 to 0.95) of the expected latent trajectory (i.e., 0.50 centile) across development and serve as the reference framework for detecting individual deviations. Notably, the latent dimensions vary in their developmental trajectories: some (e.g., latent 6 and 7) exhibit strong monotonic age-related trends, likely reflecting global structural maturation, while others (e.g., latent 2, 5, and 10) show more complex, nonlinear patterns that may capture region-specific or age-sensitive changes. This diversity suggests that the latent space encodes multiple, distinct axes of neurodevelopmental variation.

**Figure 1:**
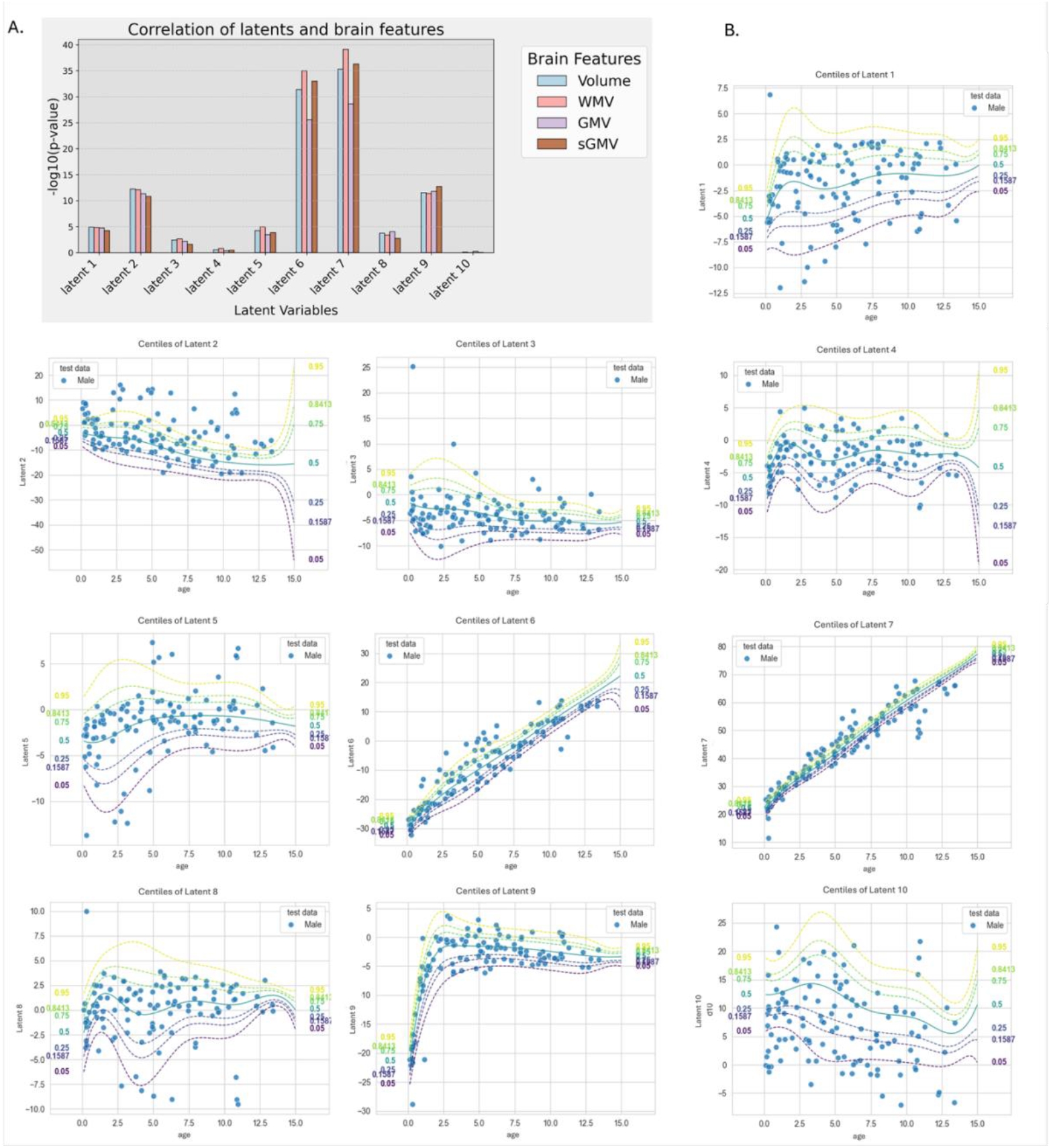
A) Bar plot showing the −log(P-value) of the correlation of each latent with brain features, including WMV), GMV, sGMV), and total brain volume. B) Normative model of latent space associated with age, serving as a reference for identifying deviations in individual brain structures. The blue dots show out of sample male participants.

We assessed the performance of the hierarchical Bayesian regression model for each latent variable using standard metrics, including mean standardized log loss (MSLL), R^2^, root mean squared error (RMSE), and correlation between predicted and observed values. Several latents (e.g., latent2, 6, 7) showed strong model fit, while others had weaker performance, likely reflecting less structured variation or deviation from the Gaussian assumption. See full results in Supplementary Table S1.

We further analysed how deviations from the normative model in the latent space (latent index) related to behavioural and cognitive functioning by testing associations with 139 measures using multiple regression. Figure 2.A presents the results of association of latent indices and cognitive/behavioural scores. To contextualise these findings, we also examined the associations between each individual brain feature and the same set of behavioural and cognitive scores. Figure 2B shows the results for sGMV, which had the strongest correlations with behavioural and cognitive measures among brain features. See Figure S2 for the corresponding analyses for WMV, GMV and total brain volume.

**Figure 2:**
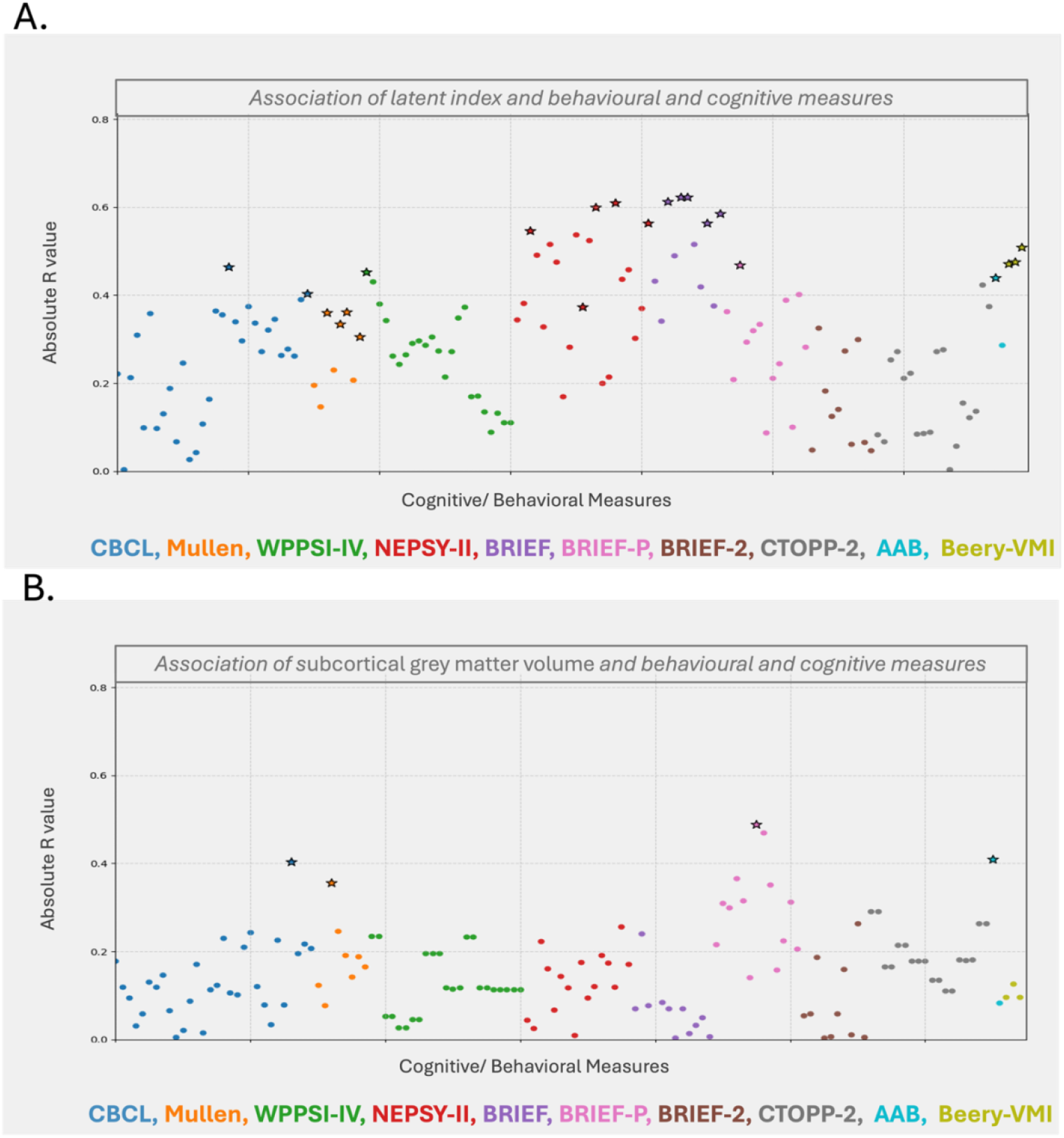
A) Manhattan plot showing the Pearson’s R values from multiple regression analyses between the latent index (deviation from the normative latent space) and 139 behavioural and cognitive measures. B) Manhattan plot showing the corresponding associations between brain features and the same set of behavioural and cognitive measures. Stars indicate significant correlations after FDR correction (∝= .05). Abbreviations: Child Behaviour Checklist (CBCL), Mullen Scales of Early Learning, Wechsler Preschool and Primary Scale of Intelligence (WPPSI-IV), Developmental Neuropsychological Assessment(NEPSY-II), Behaviour Rating Inventory of Executive Function (BRIEF), Behaviour Rating Inventory of Executive Function Preschool (BRIEF-P), Behaviour Rating Inventory of Executive Function2 (BRIEF-2), Comprehensive Test of Phonological Processing (CTOPP-2), Beery-Buktenica Developmental Test of Visual-Motor Integration (Beery-VMI), Academic Achievement Battery (AAB)

Finally, we back-projected the predicted normative latent vectors into the original input space using the decoder of the trained autoencoder. For each age and sex group, we used the posterior mean from the hierarchical Bayesian model (i.e., the expected 10-dimensional latent vector at the 50th centile) and decoded it to generate normative brain reconstructions. These synthetic scans represent the model’s estimate of typical brain structure at specific developmental stages. Figure 3.A represents ‘typical’ brain scan for male and the structural differences between males and females over time. Between 4 and 54 months, intensity increases mostly in white matter tracts, particularly in the corpus callosum and periventricular regions, potentially relating to a period of rapid early myelination. Simultaneously, cortical grey matter regions show intensity decreases. Between 54 and 108 months, white matter intensity increases become more widespread, extending to subcortical structures and frontal-parietal white matter. In contrast, we observed that cortical intensity reduces more in the frontal and temporal lobes. Finally, between 108 and 162 months, the increases in intensity persist in deep white matter and cerebellar structures, while widespread cortical reductions were observed in the frontal, temporal, and parietal cortices.

**Figure 3:**
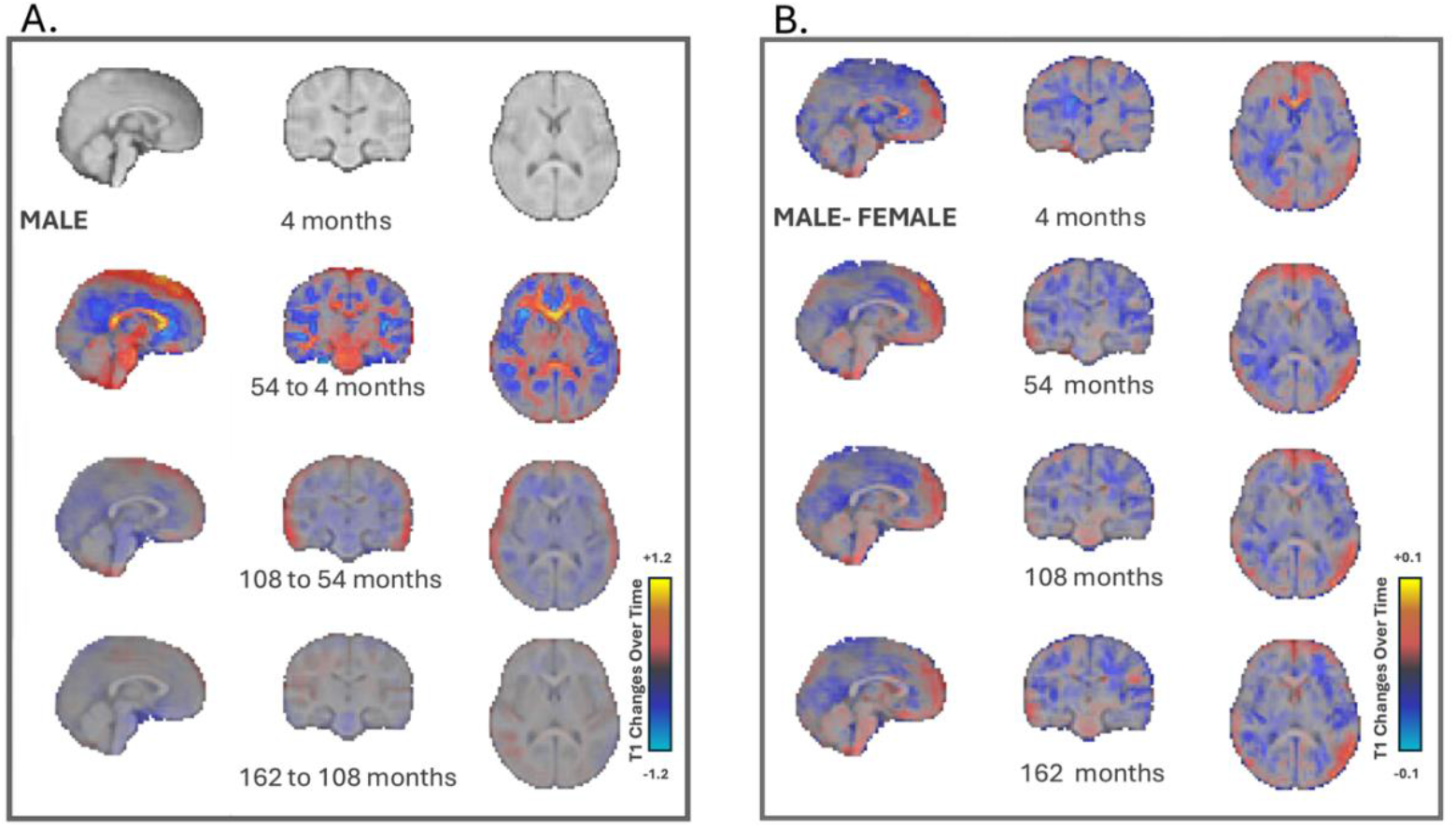
Normative latent projections mapped back to the input space. (A) Age-specific normative male brain scans reconstructed from the latent space. Brain scans demonstrate changes in intensity over time, starting at 4 months, with subsequent scans illustrating changes from 54 to 4 months, 108 to 54 months, and 162 to 108 months, capturing developmental dynamics from early infancy to adolescence. These scans illustrate ‘typical’ brain structures, aiding in the identification of deviations associated with neurodevelopmental conditions. (B) Sex-related differences in brain structure across different ages, highlighting subtle anatomical variations between males and females.

Figure 3.B shows sex-related differences in brain structure across developmental stages, derived by subtracting decoded normative scans for females from those for males at each age. Compared to the age-related changes shown in Figure 3.A, these sex differences are more subtle and spatially diffuse. Some minor regional variations can be observed at earlier ages, particularly in frontal and subcortical areas, but no consistent or large-scale patterns are apparent. At 108 months, slight male–female differences appear to emerge more distinctly in cortical regions, with males showing somewhat greater intensity in frontal and temporal white matter. These patterns persist at 162 months, with males generally exhibiting small increases in white matter intensity and reductions in cortical areas, although the effect sizes remain modest. Overall, these findings suggest that sex-related structural differences in early development are present but relatively minor compared to age-related changes.

Figure 4 shows the centiles of variation in the latent space and their corresponding projections back into the input brain space at specific age (here at the age of 4 month, see supplementary Figure S3 for age 54,108 and 162 months). The centiles represent different degrees of deviation from the normative latent representation, with the 0.5 centile reflecting the normative model. Higher centiles (e.g., 0.95) and lower centiles (e.g., 0.05) indicate increasing deviation in opposite directions from this normative baseline (0.5).

**Figure 4:**
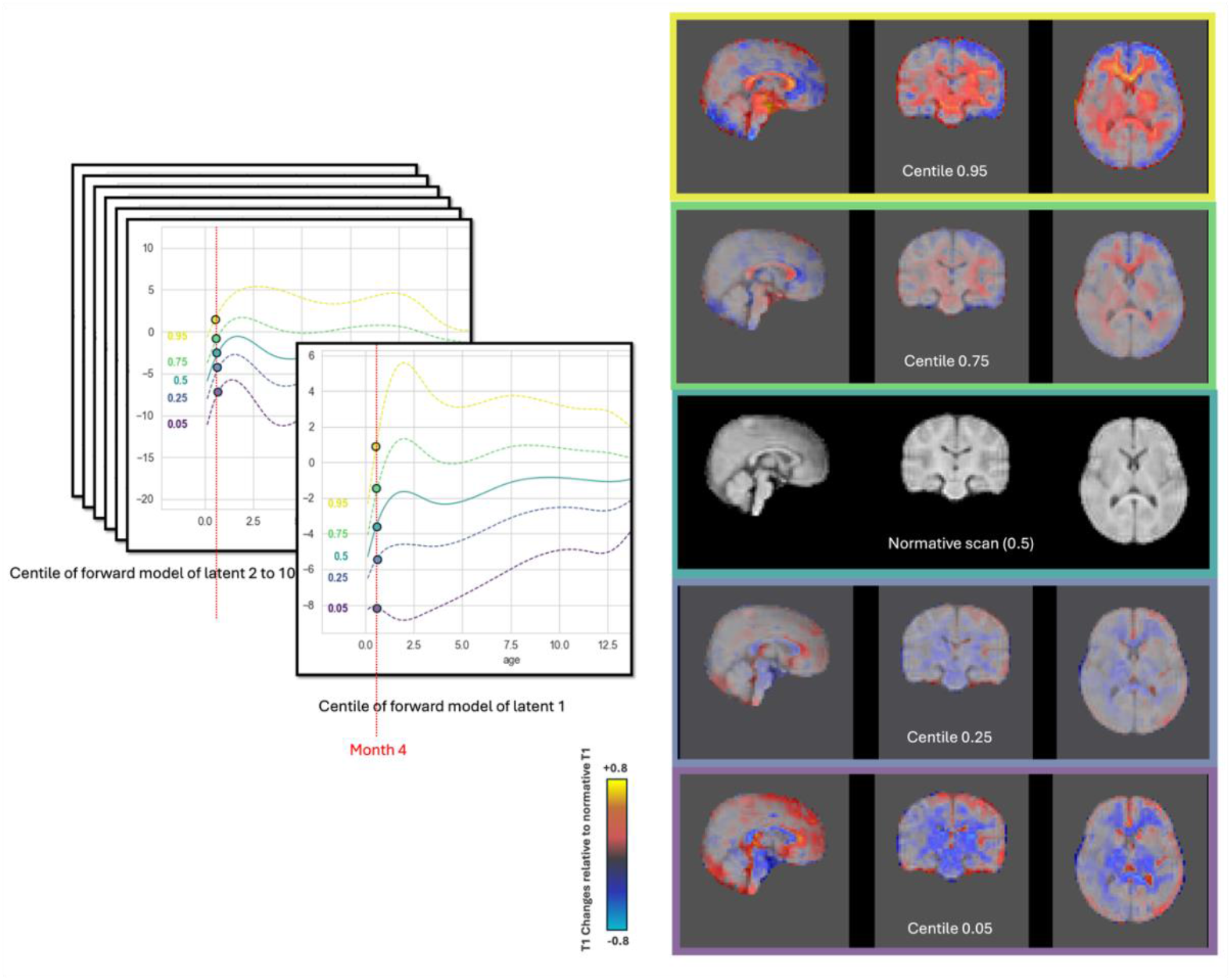
Centiles of variation in the latent space and their corresponding projections into the input space. The centre image represents the decoded brain scan from the normative latent vector (50th centile), corresponding to the model’s estimate of typical brain structure for that age and sex, while the other images show deviations at different centiles (0.95, 0.75, 0.25, and 0.05). These scans represent intensity difference maps relative to the normative scan, where higher deviations correspond to more pronounced structural differences.

## Discussion

We presented a conditional autoencoder model that highlights the potential of using latent-space multivariate normative modelling of brain-MRI data to study neurodevelopment. Our approach not only identifies significant features in neuroimaging data but also incorporates demographic variability, such as age and sex, which are essential for understanding brain development.

Our autoencoder demonstrated its efficiency through low image reconstruction error and its sensitivity to demographic variables. Specifically, the model’s ability to predict both sex and age with high accuracy indicates that it effectively learned relevant latent representations of key demographic influences on brain structure. Additionally, we were able to validate these latent representations by showing strong associations with measures of brain tissue volume. This is relevant given that brain volumes have been linked to cognitive and behavioural outcomes [38–43], though normative modelling should provide greater sensitivity then conventional volumetric measures alone.

Multiple regression analysis showed some significant relationships between deviations from the normative latent space (latent index) and behavioural and cognitive measures. Notably, the strongest associations were observed with scores from the NEPSY-II (Word Generation and Localization subtests), the CBCL Attention Problems scale, the AAB Lower Reading scaled score, and the BRIEF Metacognition Index. These findings provide external validity to the latent normative measures and suggest that deviations from the normative brain representation may reflect variability in cognition and behaviour, capturing individual differences in neurodevelopment and potentially even neurodevelopmental anomalies. An exciting possible future clinical application for this approach, could be in precision medicine, particularly psychiatry, where identifying early markers for conditions such as ASD can enable early intervention [44]. However, it is important to emphasize that these correlations require further validation in larger, more diverse populations to confirm their robustness and generalizability.

The normative brain scans generated by back-projecting the normative latent vector into the original input space offer a visual ‘benchmark’ for typical brain structures across development. While these voxel intensities do not correspond to raw MRI signal, they represent model-derived predictions of structural variation learned from the data and conditioned on age and sex. Between 4 and 54 months, the model-predicted scans show increasing voxel intensities in white matter tracts, particularly the corpus callosum and periventricular regions, potentially reflecting patterns consistent with early myelination. This developmental process improves neural transmission efficiency and is critical for the emergence of cognitive and motor functions in early childhood [45]. Concurrently, decreases in intensity within cortical grey matter regions suggest synaptic pruning, a process that refines neural circuits by eliminating redundant synapses, hence, optimizing brain function [46,47]. From 54 to 108 months, the widespread increase in white matter intensity extending to subcortical structures and frontal-parietal regions reflects ongoing myelination, supporting the maturation of complex cognitive functions. The observed cortical intensity reductions, particularly in the frontal and temporal lobes, align with cortical thinning associated with synaptic pruning and grey matter reduction, processes that continue into adolescence to enhance neural efficiency[3,48].

From 108 to 162 months, persistent intensity increases in deep white matter and cerebellar structures indicate continued myelination, essential for the refinement of motor and cognitive skills. Simultaneously, widespread cortical reductions in the frontal, temporal, and parietal cortices correspond with ongoing synaptic pruning and cortical thinning, processes that are vital for the development of higher-order cognitive functions during adolescence. These developmental trends align with previous studies on the patterns of white matter expansion and cortical maturation throughout childhood and adolescence [45,47,49–55].

While the developmental changes across sex are not pronounced in comparison, they align with previous findings indicating that males experience more prominent white matter maturation during development [56,57]. These patterns are consistent with findings that males undergo more significant age-related reductions in grey matter while maintaining greater increases in white matter volume, potentially influencing cognitive and behavioural trajectories differently across sexes [42,54,55,58,59].

The centile plots in Figure 4 illustrate how deviations in the latent space relate to structural changes in the generated brain scans. Brain images generated from latent vectors near the normative centre (e.g., the 0.5 centile) show minimal deviation from the normative scan, reflecting typical structure. As we move away from the centre, toward either lower (e.g., 0.05) or higher (e.g., 0.95) centiles. the reconstructed images exhibit increasingly pronounced structural differences. Notably, these changes appear continuous and directionally consistent, with deviations on either side of the normative latent vector producing mirrored patterns. This suggests that the latent space captures meaningful axes of variation in brain structure, with movement along these axes corresponding to systematic, interpretable changes in the reconstructed anatomy. These scans not only provide a useful reference for identifying deviations in individual brain structure but also serve as a potential clinical tool for researchers and clinicians to assess structural differences associated with neurodevelopmental disorders. By comparing individual brain scans to these normative baselines, clinicians would be provided with 3D maps of neurodevelopmental deviations that could signal health concerns, which could contribute to a more tailored approach in both diagnosis and treatment planning.

Our study has some limitations. Given the relatively small sample size, the autoencoder is more likely to capture broad intensity-based patterns distributed across the brain, rather than localized or sharply defined morphological features such as cortical thickness or shape deformations. As a result, the decoded normative scans primarily reflect smooth structural variation learned from voxel-wise signal patterns, rather than detailed anatomical differences. A larger, more diverse dataset would enable the model to capture more variability, such as differences in brain shape, and potentially improve its ability to model complex neurodevelopmental trajectories. Furthermore, although our study is cross-sectional, incorporating longitudinal data would significantly enhance our ability to track and predict dynamic changes in brain structure over time. This is particularly important in neurodevelopmental research, where understanding the trajectory of brain growth and how it aligns with behavioural milestones can provide more predictive power for cognitive and behavioural outcomes.

Another limitation relates to the distributional assumptions in the normative modelling step. We assumed a Gaussian likelihood for the normative modelling of latent variables, as the autoencoder was trained using mean squared error (MSE), which implicitly encourages normally distributed residuals. Additionally, the data preprocessing (z-scoring) and architectural choices (e.g., L1 regularization, bias removal) promote symmetry and sparsity in the latent space, supporting the assumption of approximate normality. While this assumption holds for several latents with strong demographic structure, future work may benefit from more flexible likelihoods to better capture non-Gaussian dimensions.

Future research should focus on expanding the dataset to include larger and more diverse samples, allowing for a broader representation of individual variations in brain structure, particularly from underrepresented populations, based on ethnicity and geography [60,61]. In addition, future studies could integrate multimodal neuroimaging, combining more structural MRI modalities (e.g., T2w-MRI, DWI) alongside functional data such as fMRI. This would provide a more comprehensive understanding of how both structural and functional brain changes contribute to behaviour and cognition.

In conclusion, our study demonstrates that a 3D semi-supervised autoencoder provides a framework for capturing individual variations in brain development through a multivariate normative model. By leveraging latent representations of paediatric MRI data, our approach identifies structural patterns associated with age and sex while preserving both local and global features of brain morphology. The ability to generate normative brain scans from the latent space further enhances interpretability, offering a benchmark for understanding typical neurodevelopmental trajectories. By integrating additional demographic, behavioural, and multimodal data, this method holds a potential for advancing our understanding of brain-behaviour relationships and supporting precision medicine approaches in neurodevelopmental research.

## Supporting information

Supplementary docs

