## Supplementary docs for "Latent Representations of Early Brain Development: A Multivariate Normative Model of Brain Structure and Behaviour"

**Supplementary materials**

### Methods

Figure S1 shows the methods overview.

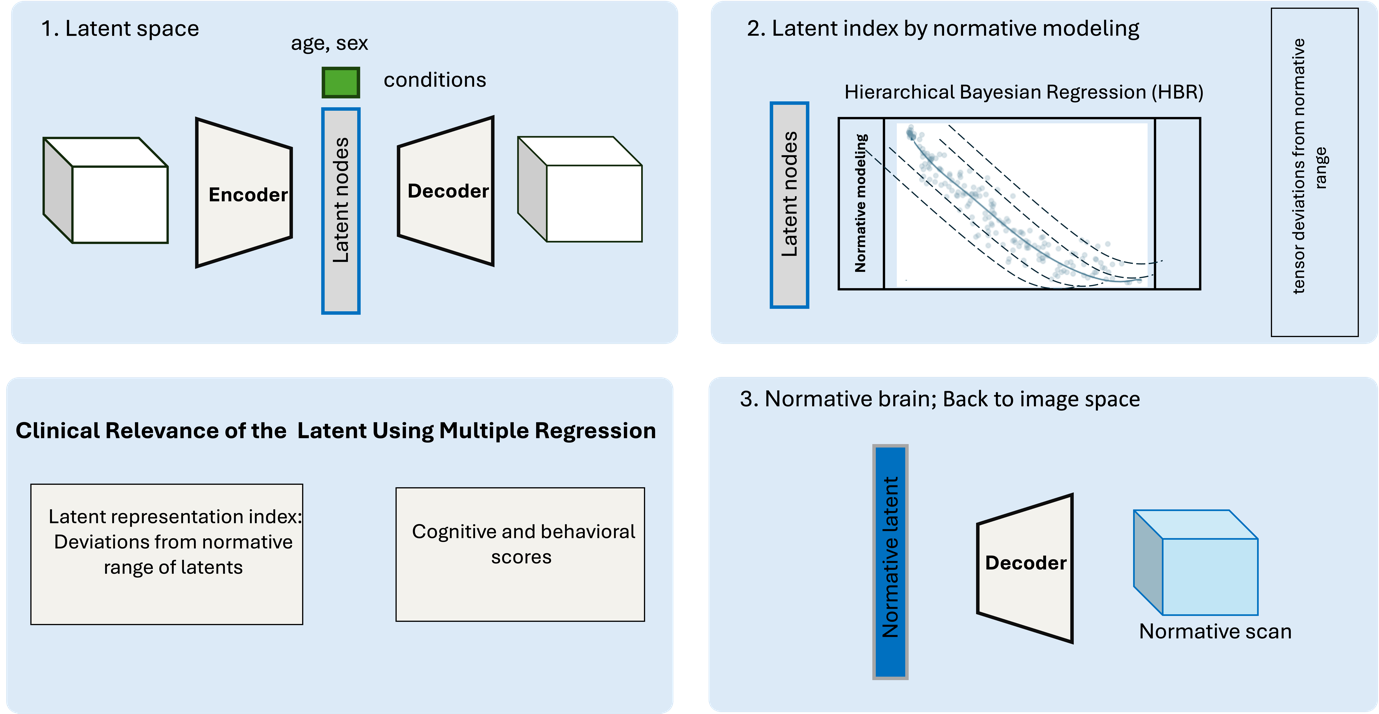

### Autoencoder Architecture

We implemented a 3D conditional autoencoder with demographic prediction capabilities. The encoder consisted of three convolutional layers (64, 32, 16 filters) with 3×3×3 kernels, LeakyReLU activations, average pooling, batch normalization, and dropout (0.1). The network compressed input images to a 10-dimensional latent space while simultaneously predicting age and sex through dedicated output branches. The decoder mirrored this structure with three transposed convolutional layers (16, 32, 128 filters), upsampling operations, and a final reconstruction layer. Both networks were implemented without bias terms to reduce model complexity.

### Training Procedure

The model was trained using a composite loss function:

L=Reconstruction Loss+ age prediction Loss+ sex prediction Loss+⋅αL1+⋅βL2

where Reconstruction Loss is the mean squared error between input and reconstructed images (with additional weighting for masked regions), age prediction Loss is the mean absolute error for age prediction, sex prediction Loss is binary cross-entropy for sex classification, and L1 and L2 are regularization terms (α=e-3+⋅β=e-2). We employed the Adam optimizer (learning rate = e-3, weight decay = e-4) with gradient clipping (max norm = 1.0). Training proceeded for 1,000 epochs with batch size 32, reserving 8% of training data for validation.

Normative modelling analysis was done using pcntoolkit (https://pcntoolkit.readthedocs.io/en/latest/index.html)

### Results

Table S1 reports model fit metrics for each latent variable using hierarchical Bayesian regression with a Gaussian likelihood. Metrics include mean standardized log loss (MSLL), negative log likelihood (NLL), coefficient of determination (R²), root mean square error (RMSE), Pearson correlation (ρ), p-value of the correlation, standardized mean squared error (SMSE), and the Shapiro-Wilk statistic assessing normality of the residuals.

| **Latent** | **MSLL** | **NLL** | **R2** | **RMSE** | **Rho** | **Rho_p** | **SMSE** | **ShapiroW** |
| --- | --- | --- | --- | --- | --- | --- | --- | --- |
| latent1 | 1.0540 | 1.5492 | 0.0909 | 3.1164 | 0.3063 | 0.0000 | 0.0219 | 0.9069 |
| latent2 | 2.8506 | 0.8997 | 0.6572 | 6.0271 | 0.8243 | 0.0000 | 0.0022 | 0.9519 |
| latent3 | 1.4013 | 1.4370 | 0.0812 | 3.9629 | 0.2953 | 0.0000 | 0.0048 | 0.8874 |
| latent4 | 1.3788 | 1.3462 | 0.0755 | 3.5499 | 0.3148 | 0.0000 | 0.0133 | 0.9556 |
| latent5 | -5.7578 | 8.4487 | 0.1129 | 3.3608 | 0.3642 | 0.0000 | 0.0110 | 0.4260 |
| latent6 | 3.7652 | 0.1894 | 0.9223 | 3.5211 | 0.9590 | 0.0000 | 0.0005 | 0.9156 |
| latent7 | 0.5519 | 3.4570 | 0.9721 | 2.2264 | 0.9880 | 0.0000 | 0.0001 | 0.5409 |
| latent8 | -0.3973 | 2.9455 | 0.0932 | 2.9459 | 0.3782 | 0.0000 | 0.0147 | 0.7356 |
| latent9 | -99.7419 | 103.0147 | 0.8137 | 2.7560 | 0.6804 | 0.0000 | 0.0015 | 0.0745 |
| latent10 | 2.7328 | 1.0127 | 0.6204 | 6.3114 | 0.7563 | 0.0000 | 0.0028 | 0.9219 |

Figure S2 shows the associations between individual brain features white matter volume (WMV), grey matter volume (GMV), subcortical grey matter volume (sGMV), and total brain volume, and the behavioural and cognitive measures.

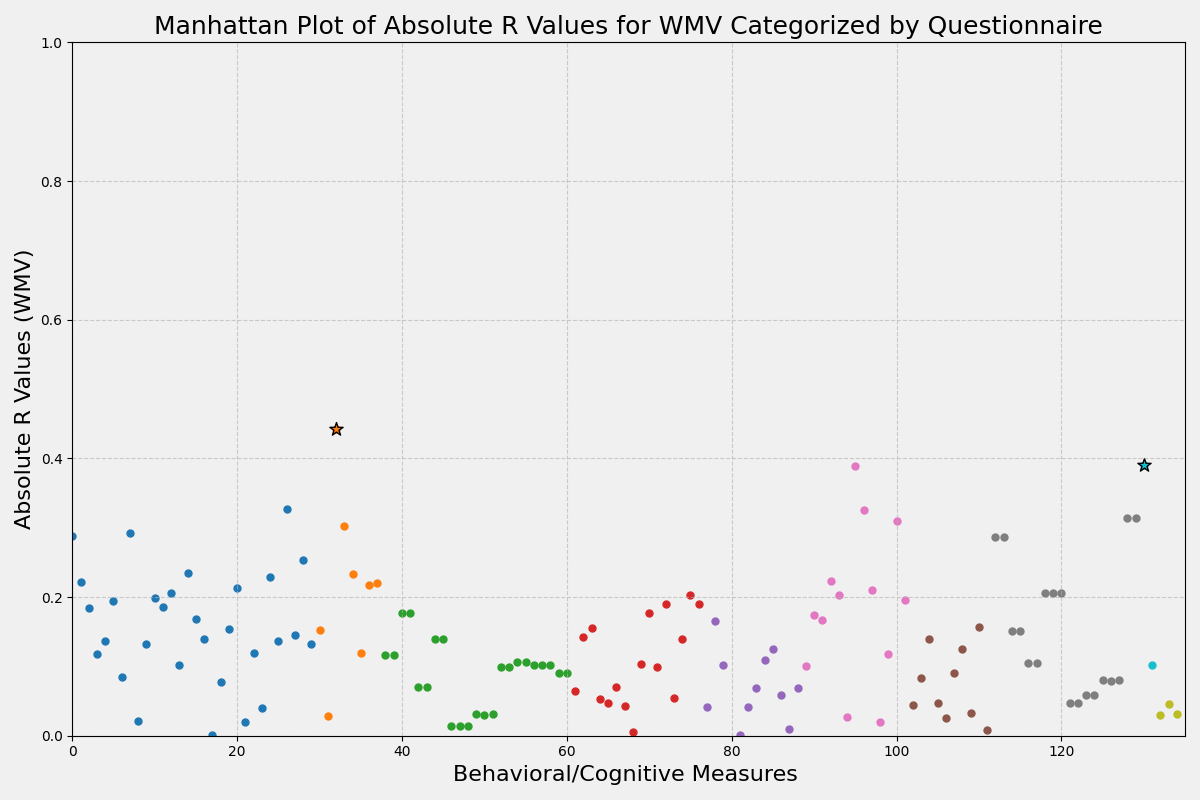

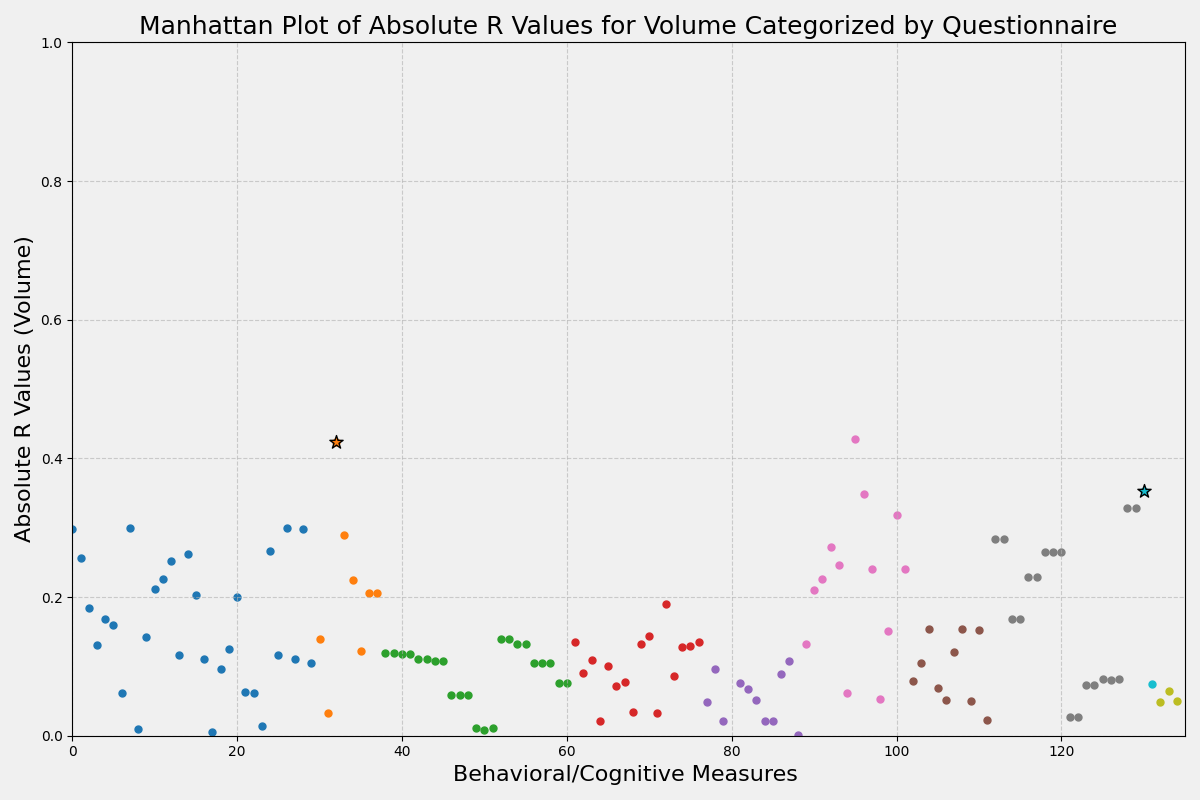

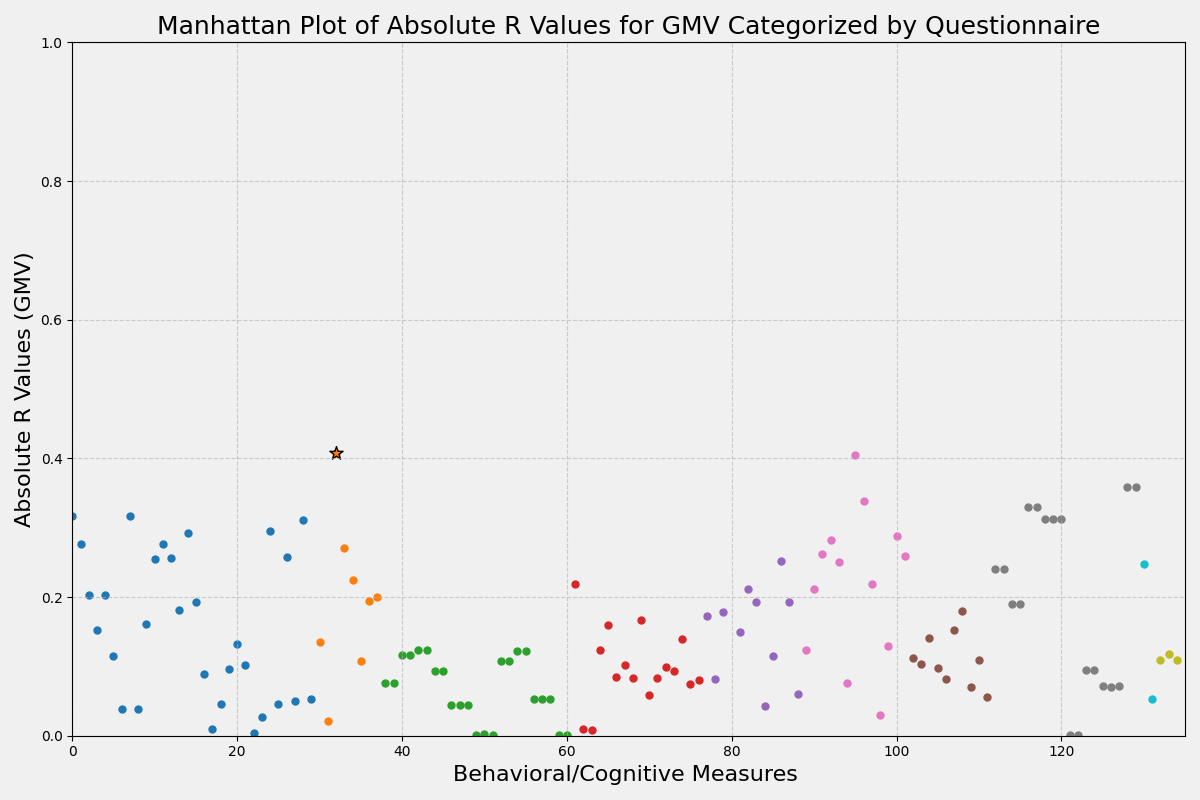

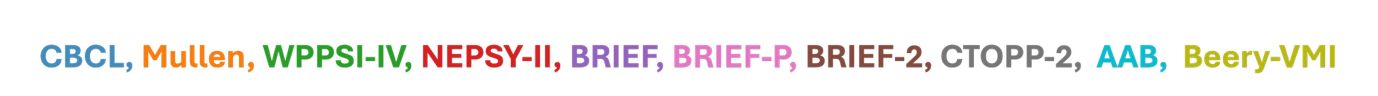

Figure S2: Associations between individual brain features (WMV, GMV, sGMV, and total brain volume) and behavioural and cognitive measures. Stars indicate significant correlations after FDR correction ($\propto=.05$). Abbreviations: Child Behaviour Checklist (CBCL), Mullen Scales of Early Learning, Wechsler Preschool and Primary Scale of Intelligence (WPPSI-IV), Developmental Neuropsychological Assessment(NEPSY-II ), Behaviour Rating Inventory of Executive Function (BRIEF), Behaviour Rating Inventory of Executive Function Preschool (BRIEF-P), Behaviour Rating Inventory of Executive Function 2 (BRIEF-2), Comprehensive Test of Phonological Processing (CTOPP-2), Beery-Buktenica Developmental Test of Visual-Motor Integration (Beery-VMI), Academic Achievement Battery (AAB)

Figure S3 shows the centile of variation at the age of 54,108 and 162 months, respectively.

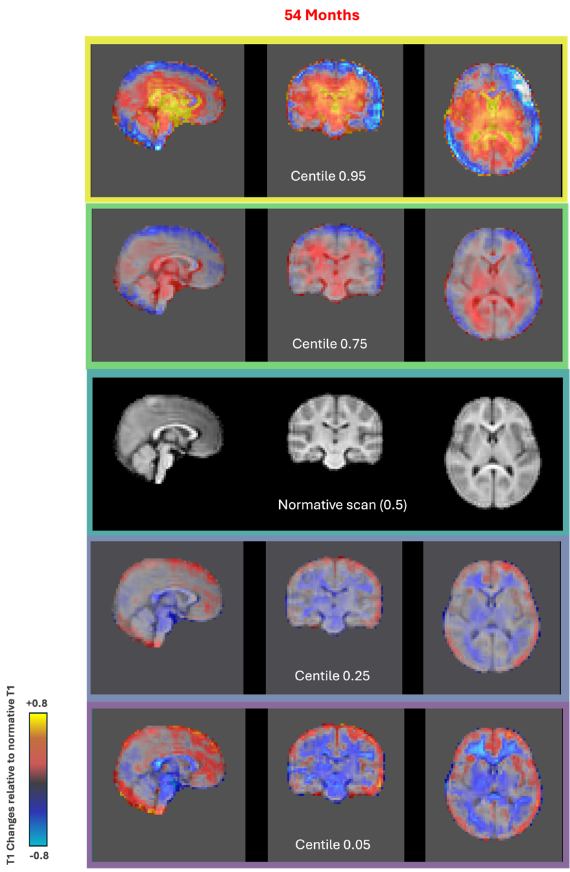

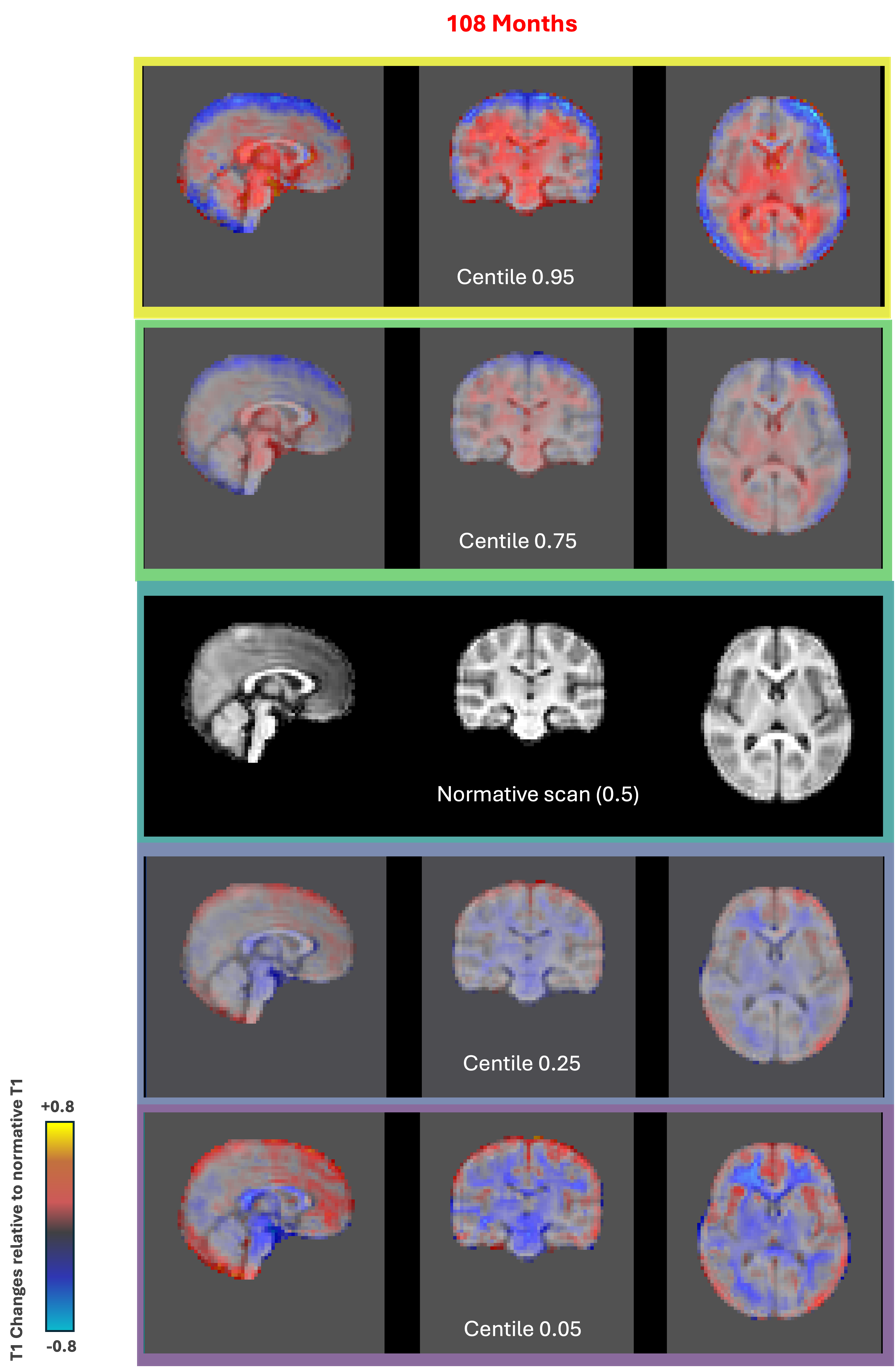

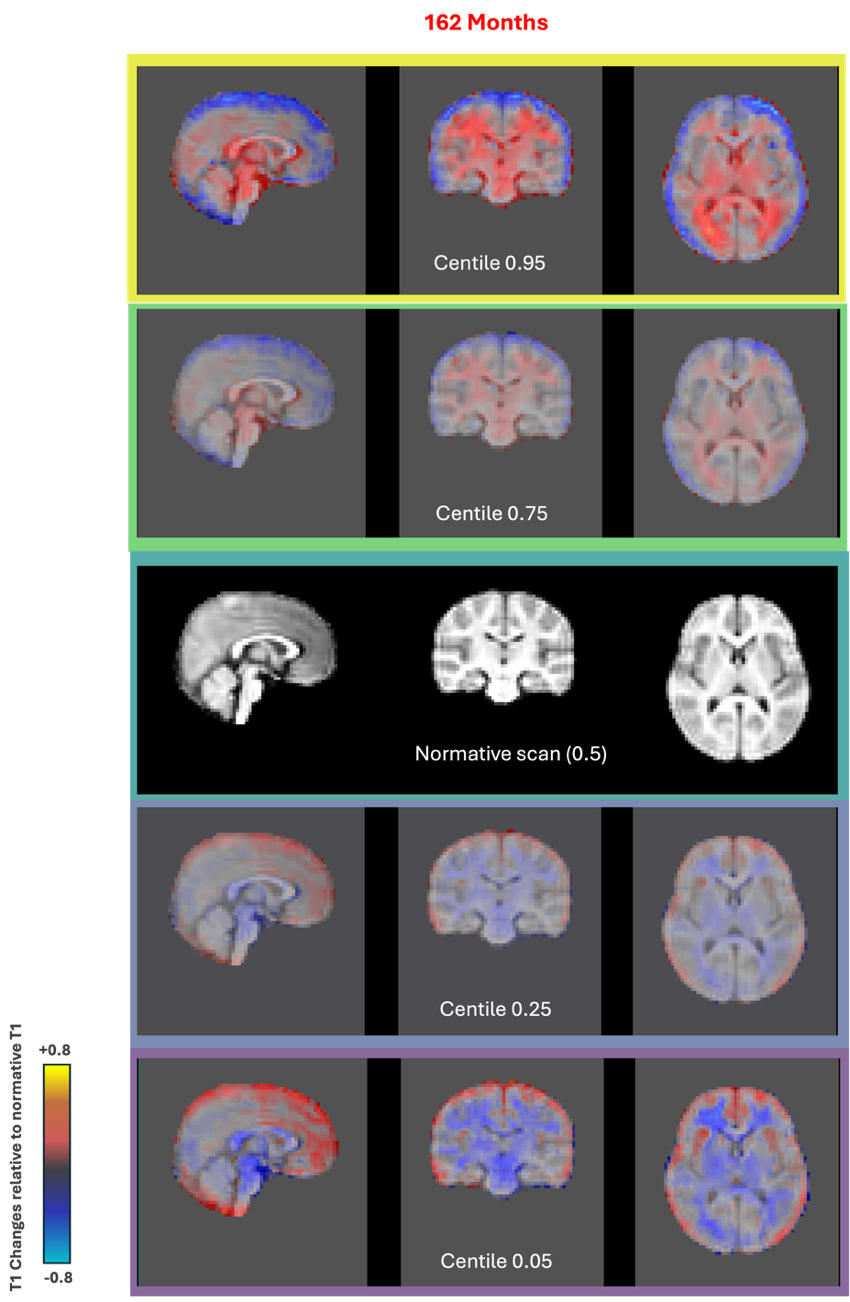

Figure S3: Centiles of variation in the latent space and their corresponding projections into the input space. The normative scan (centre) represents the typical brain structure, while the other images show deviations at different centiles (0.95, 0.75, 0.25, and 0.05). These scans represent intensity difference maps relative to the normative scan, where higher deviations correspond to more pronounced structural differences.
